# Single pointwise samples of electric field on a neuron model cannot predict activation threshold by brain stimulation

**DOI:** 10.1101/2025.07.31.667968

**Authors:** Boshuo Wang, Minhaj A. Hussain, Torge Worbs, Axel Thielscher, Warren M. Grill, Angel V. Peterchev

## Abstract

**Background:** Some computational models of neural activation by transcranial magnetic stimulation overestimate the electric field (E-field) threshold compared to *in vivo* measurements. A recent study proposed a statistical method to account for the influence of microscopic perturbations to the E-field. The method, however, relies on the unsubstantiated assumption that thresholds can be predicted by single pointwise samples of the E-field strength along neural cables.

**Objective:** We analyzed neural responses to E-field with microscopic perturbations and demonstrate via theoretical derivation and simulations that neural activation is not determined by pointwise E-field amplitude but rather by spatial integration of the E-field along the neural cable. Therefore, the influence of microscopic E-field perturbations is negligible due to the spatiotemporal filtering by the neural membrane and axoplasm.

**Methods:** We derive the axial and transmembrane currents in a neural cable for an imposed E-field with microscopic perturbations. We simulate neural activation thresholds of unmyelinated and myelinated axons in two stimulation scenarios and compare thresholds for E-field activation with and without perturbations.

**Results:** In the theoretical derivation, the perturbation terms average out to zero on larger spatial scales indicating that they do not influence neural activation thresholds. Simulated thresholds with the E-field spatial perturbations present had negligible differences (< 3.4%) compared to those without.

**Conclusion:** Single point samples of the microscopic E-field on a neural cable cannot predict neural activation thresholds. Neural simulations should be used to determine any influence of the E-field spatial perturbations. The latter, however, are unlikely to account for the difference between experimental and simulated E-field thresholds.

## Introduction

Computational modeling can help uncover the neural mechanisms of transcranial magnetic stimulation (TMS). However, electric field (E-field) thresholds simulated using morphologically-realistic neurons embedded in head models [1,2] were higher than those from human experiments [3], and resolving this mismatch is an important effort. Weise et al. attempted to reconcile this threshold discrepancy using a statistical model [4] based on the E-fields calculated from microscopic tissue structures^1^[5] to account for E-field perturbations due to interactions with dense cellular membranes. The perturbations have zero spatial mean but yield many local microscopic “hotspots” with substantially higher amplitude than the average E-field. Using the microscopic E-field sampled along axonal segments and assuming that neural activation occurs if the local E-field exceeds a threshold amplitude, Weise et al. derived a recruitment function that generated neural activation behavior matching experimental TMS [4].

The model [4], however, relied on the assumption that activation thresholds of a model neuron can be determined by pointwise samples of a highly nonuniform E-field from a single location along the neural cable. This implies the recruitment behavior of the single pointwise E-field sample from an arbitrary field distribution is the same as a uniform E-field of the same amplitude spanning the entire neuron. Specifically, Weise et al. considered axon terminals, which in some models have the lowest activation thresholds within the axonal arbor [1,2], to be activated if the local microscopic E-field exceeded the threshold of the whole cell under a uniform E-field. This assumption markedly oversimplifies the behavior of neural activation and may limit the validity of the results.

### Theoretical framework

The E-field generated by non-invasive brain stimulation is often considered uniform on the scale of individual neurons [6], and thresholds are often reported as E-field amplitudes that are macroscopic averages of the microscopic field [7]. Exogenous E-fields drive neuronal polarization according to the co-axial E-field along the elements, with the activating function being proportional to E-field strength at terminals and E-field derivative mid-cable [8]. In a uniform E-field, mid-cable activation is determined only by the curvature and changes in diameter of the axon. When microscopic E-field perturbations are present, the activating function can appear to increase dramatically, i.e., the E-field derivative can be very large. However, due to the high conductivity of the axoplasm compared to the membrane, transmembrane potentials from nearby locations are tightly coupled via axial ohmic current flow, and thus neural activation is determined by the complex interaction of the external activating function, internal current redistribution, and local membrane dynamics over time [9].

We consider the total E-field consisting of the macroscopic (average) field and the local microscopic field due to local tissue inhomogeneities:

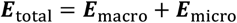

Here, ***E***_micro_ has zero mean^2^on spatial scales larger than the local inhomogeneities, i.e., the diameter of somas, neurites, and blood vessels [5]. In the discretized neural cable (Fig. 1), the axial intracellular current *I*_i_ is given by

**Figure 1.**
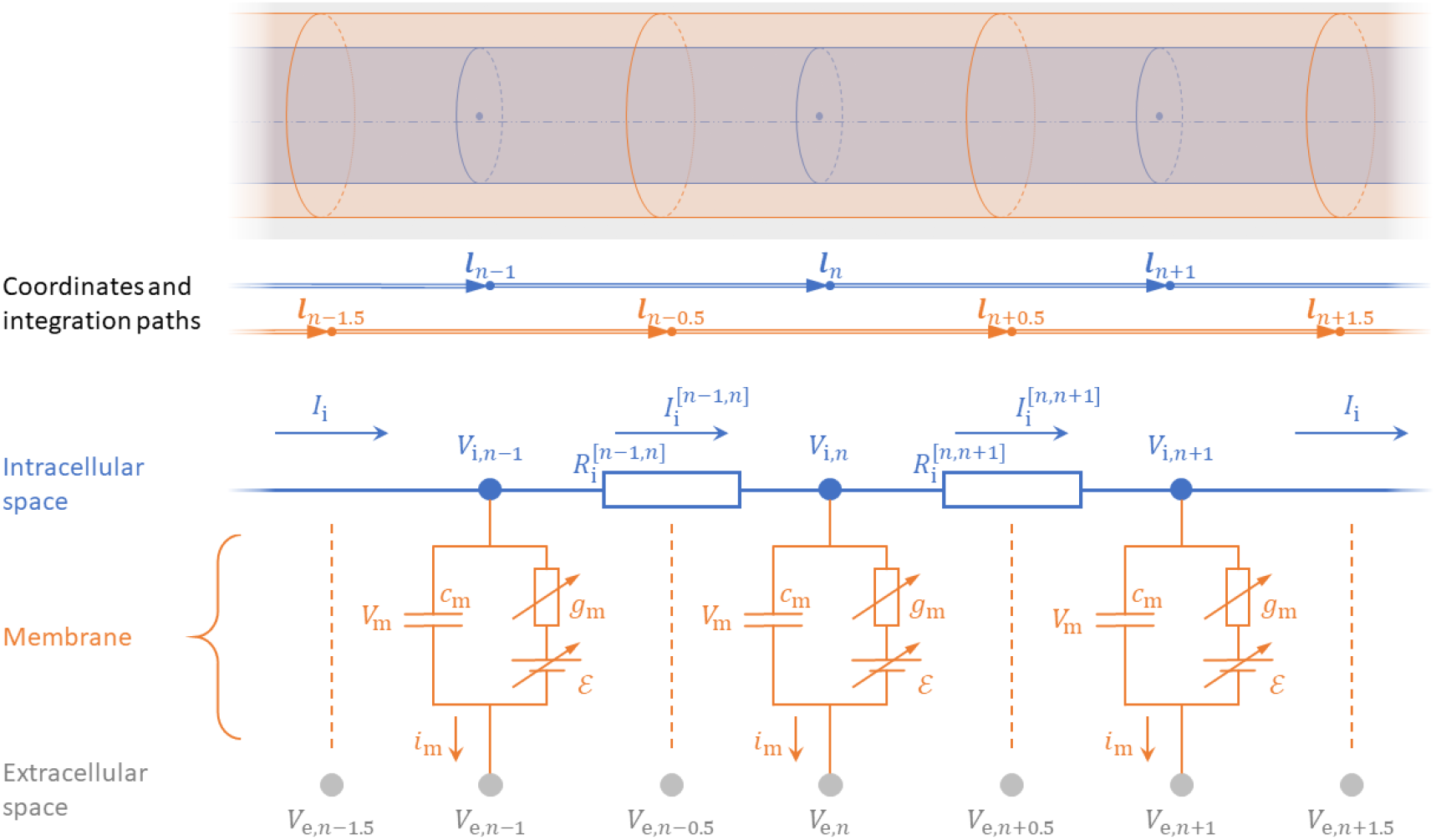
Neural cable model showing the discretized axoplasm (blue) and membrane (orange). The compartment centers (blue dots, at coordinates with integer subscripts), which define the intracellular potentials, are connected by axial resistance determined by the distances between the centers. The neural membranes, which define the transmembrane potentials, span the distances between compartment boundaries (at coordinates with subscripts offset by 0.5). The integration paths for axial and transmembrane currents are therefore staggered, with the coordinates potentially with non-even spacing.

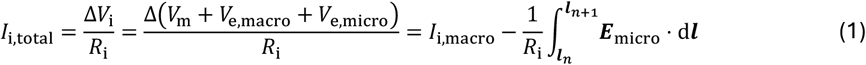

Here, *R*_i_ is the axial resistance between compartments, Δ*V*_i_ is the intracellular potential^3^difference, the transmembrane potential is *V*_m_ = *V*_i_ − *V*_e,total_, and the lower and upper bounds of the integral, ***l***_*n*_ and ***l***_*n*+1_, are respectively the center coordinates of the *n*-th and (*n* + 1)-th compartments. Considering reduced potentials, the transmembrane current *i*_m_ for a given compartment is

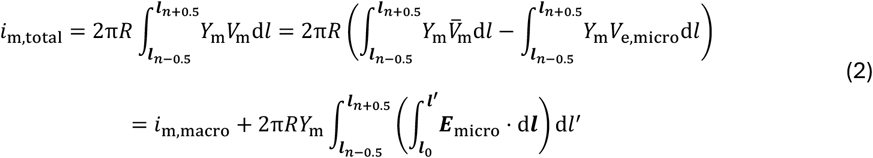

Here, *R* is the compartment radius; *Y*_m_ = *g*_m_ + *j*ω*c*_m_ is the specific membrane admittance in the frequency (ω) domain, with *g*_m_ and *c*_m_ respectively being specific membrane conductance and capacitance; 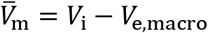 is the average transmembrane potential under macroscopic E-field only; the lower and upper bounds of the outer integral, ***l***_*n*−0.5_ and ***l***_*n*+0.5_, are respectively the coordinates of the boundaries between *n*-th compartment and its two neighbors; and ***l***_0_ is the coordinate of the reference potential (e.g., soma).

In both calculations, the microscopic terms are zero on average over the integration distances. Especially for transmembrane currents *i*_m_, the influence of the microscopic E-field is reduced further compared to that of the axial current *I*_i_, due to suppression of high spatial frequencies by the double integration. Although the microscopic E-field would appear to affect these electrical currents if the spatial discretization is chosen to be very small, the highly conductive axoplasm results in tight coupling of these small compartments, and the overall neural response is still determined by their collective behavior over the larger scale of the neural cable’s length constant. Therefore, *I*_i,total_ = *I*_i,macro_ and *i*_m,total_ = *i*_m,macro_, and the microscopic E-field does not affect the response of the cable due to spatiotemporal filtering [7], resulting in approximately the same activation threshold without the perturbations [5].

### Neural activation thresholds in the presence of E-field spatial micro-inhomogeneities

To demonstrate the minimal effect of microscopic E-field inhomogeneities, we simulated neural activation thresholds for two scenarios: a terminating axon in an aligned quasi-uniform E-field (the context for the analysis by Weise et al. [4], Fig. 2A) and placed near a spherical electrode (Fig. 2B).

**Figure 2.**
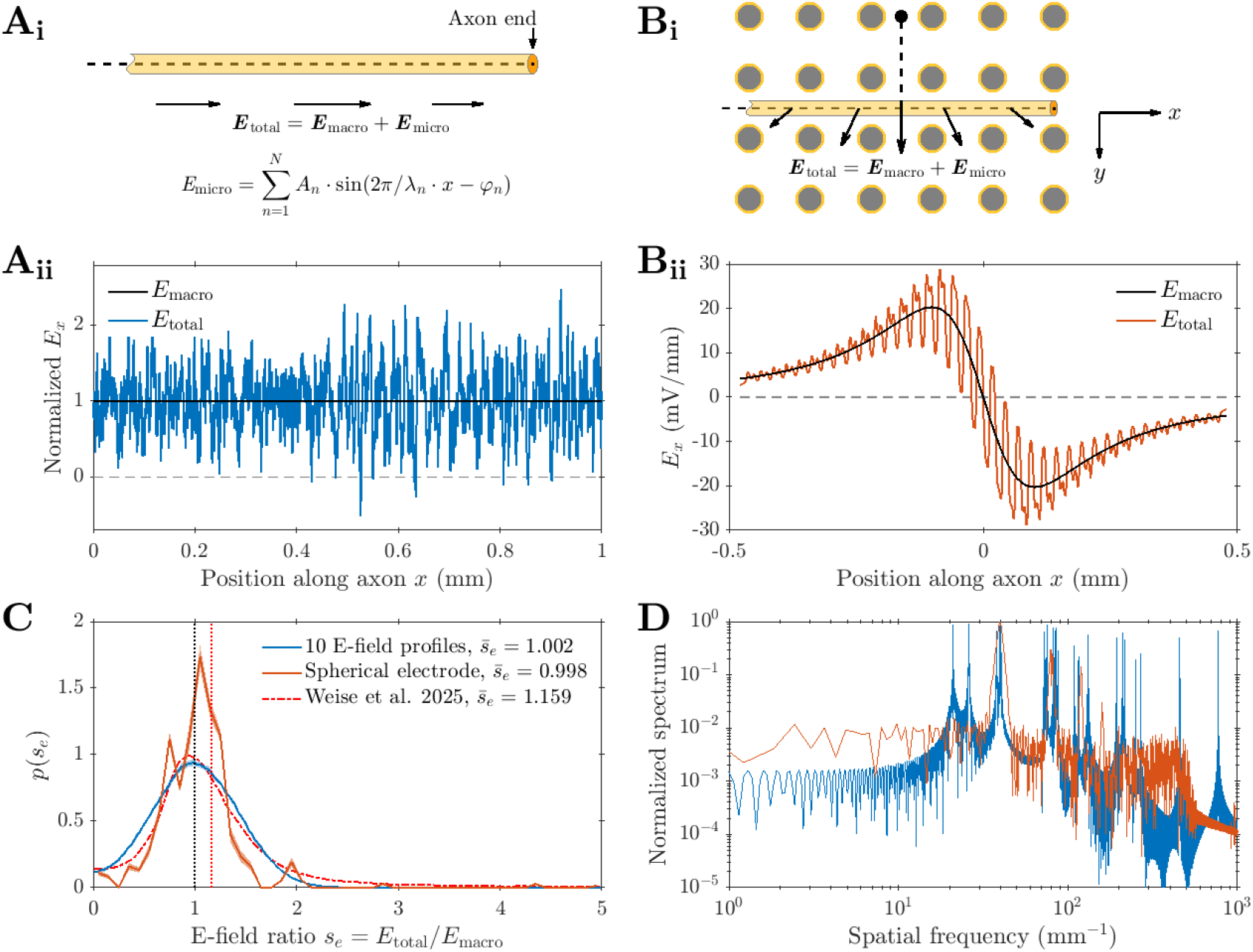
**A. i**. A uniform macroscopic E-field with sinusoidal microscopic spatial variations imposed on an aligned terminating axon (yellow). **ii**. Normalized E-field profile along the axon, showing one example out of the ten generated. **B. i**. A non-uniform macroscopic E-field from a 50 μm radius spherical electrode (black dot, anodic current) is perturbed by three interleaved grids of axons (cross-sections of axons shown as gray disks with yellow outlines; the two grids parallel to the *x* − *y* plane not shown). The target axon was modeled to be 83 μm from the electrode surface, extend beyond the original simulation space in the negative *x* direction, and terminate at 40 locations in the positive *x* direction. **ii**. E-field profile along the axon for a cathodic current of 8 μA. The macroscopic field was obtained for homogenous conductivity of 0.6 S/m to match the effective conductivity of the axon grid embedded in 1 S/m extracellular space. [7] **C**. Probability density distribution of the ratio between the total and macroscopic fields *s*_e_ = *E*_total_/*E*_macro_, with their mean indicated by the vertical dotted lines. The distribution was calculated as an average of 10 profiles for the scenario in panel B. The distribution from [4] (red dashed line) has a mean significantly larger than one, indicating that the microscopic E-field has non-zero mean. **D**. Spatial frequency spectrum of the total E-field, with one example profile for the scenario in panel A.

## Methods

For the aligned E-field, the microscopic variations in the E-field consisted of 25 sinusoids with wavelengths randomly sampled from uniform distributions between 1 and 15 μm (20 samples) and 15 and 50 μm (5 samples) to represent respectively perturbations from small (e.g., axons and dendrites) and large (e.g., blood vessels and large somas) structures^4^. The sinusoids had random phases and their amplitudes were fitted so that the distribution of E-field ratio *s*_e_ = *E*_total_/*E*_macro_ matched the study by Weise et al. [4] as closely as possible (Fig. 2C). Ten such microscopic E-field profiles were generated. For the spherical electrode, the E-field profiles from [7] were used. The E-field was sampled along a line 83 μm from the surface of a 50 μm radius electrode. The total E-field with microscopic variations was generated by three interleaved grids of axons with 4.46 μm radius and 25 μm spacing (30% volumetric fraction) in an extracellular space of 1 S/m conductivity, whereas the macroscopic field was from a homogenous space with conductivity of 0.6 S/m matching the effective conductivity of the interleaved axon grids.

Unmyelinated and myelinated axons of 0.3 and 3 μm radius (length constants of 250 μm and 790 μm for unmyelinated axons and 104 μm and 1.13 mm for myelinated axons, respectively) were simulated [8] and thresholds were determined for rectangular pulses with 7 pulse durations between 10 μs and 10 ms within 0.05% accuracy. Thresholds obtained with the total field were compared against those with the macroscopic E-field only. For the aligned E-field, the axon was 1 cm long and placed at 10 random positions along each of the 10 E-field profiles, thus yielding 100 simulations per pulse duration. For the spherical electrode, the axons were “semi-infinitely long” in the negative *x* -direction, i.e., extended beyond the original simulation space of 0.5 mm radius with their terminals deactivated, and they terminated in the positive *x*-direction at 40 uniformly spaced locations between 0 and 0.5 mm to capture both terminal and mid-axon activation. Simulation time step was 1 μs and spatial discretization was 2 μm.

## Results

The ratio between the total and macroscopic E-field, *s*_e_, for both the axons in the aligned E-field and next to the spherical electrode followed a similar distribution that peaked around 1, with the mean ratio very close to 1 (Fig. 2C). In contrast, the distribution in Weise et al. [4] had a mean of 1.16, which corresponded to a non-zero mean and was likely a result of applying insufficient scaling to the microscopic E-field to match the macroscopic field [5,7]. The spatial spectra of the E-field variations (Fig. 2D) show that the spatial frequencies correspond to wavelengths in the expected range between 1 and 50 μm, matching the spacing of the perturbations.

The relative differences between thresholds obtained with the total E-field versus those with the macroscopic E-field were small and negligible across all pulse durations (Fig. 3): for unmyelinated axons, changes were < 0.3% for 3 μm radius and < 2% for 0.3 μm radius, and for myelinated axons, threshold changes were < 1.9% for 3 μm radius and < 3.4% for 0.3 μm radius. Although strong insulation by myelin should in general further reduce the influence of microscopic E-field^5^ compared to that of unmyelinated axon [8,10], the two models here have different properties, with the myelinated axons having ion channels with higher conductance and faster dynamics and thus being more sensitive to changes in the E-field.

**Figure 3.**
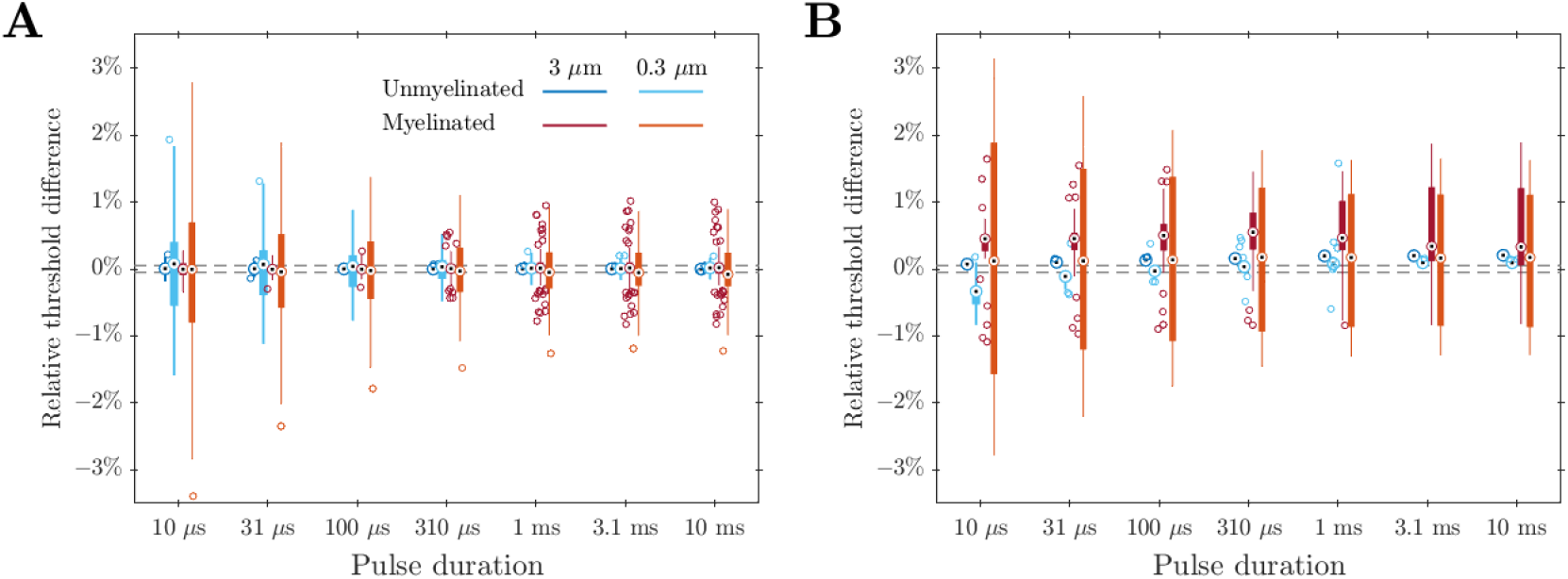
Relative differences of thresholds of unmyelinated (blue) and myelinated (red) axons [8] of 0.3 and 3 μm radius under the total E-field versus those in response to the macroscopic E-field for 7 pulse durations. **i**. Statistics of 100 thresholds per pulse duration obtained with uniform macroscopic E-field. 10 thresholds were calculated for each of the 10 E-field profiles, with the axon shifted along the profiles. **ii**. Statistics of 40 thresholds per pulse duration obtained with non-uniform macroscopic E-field from spherical electrode. The box-and-whisker plots show median (circles with center dots), first and third quartiles (boxes), upper and lower adjacent values (whiskers), and any outliers above or below the adjacent values (open circles). The dashed horizontal lines indicate ±0.05%, the relative accuracy for threshold calculation.

## Discussion and conclusions

Our analysis and simulations demonstrate that neural activation thresholds are determined by the aggregate influence of the E-field over the length of the neural cable and not the pointwise E-field amplitude. Due to the tight electrical coupling within the axoplasm, the effects of the microscopic E-field variations average out to be the same as the macroscopic field without the variations, despite the extremely fine spatial resolution used in the simulation. The E-field should be considered on the spatial scale of the cable’s length constants. Only when comparing different stimulation scenarios that have the same relative E-fields distributions along the neural cable or across the entire cell (whether (quasi-)uniform [1,2] or non-uniform [11]) and the same temporal stimulation waveform, can activation threshold be considered as inversely-proportional to the (peak) E-field amplitude and the E-field amplitude be used as a predictor of relative thresholds. Otherwise, neural simulations should be performed to solve the cable equation.

Similar misestimation of global neural behavior based on local fields occurred previously with transverse polarization, where the local transmembrane potential could deviate significantly from the average value around a neural compartment’s circumference (Fig. 4) [8]. When linear integrate-and-fire behavior is applied pointwise to local transmembrane potentials, thresholds were predicted to change significantly due to transverse polarization, but neural simulations considering the transmembrane current integrated over the entire compartment’s circumference demonstrated that this was not the case [8,10].

**Figure 4.**
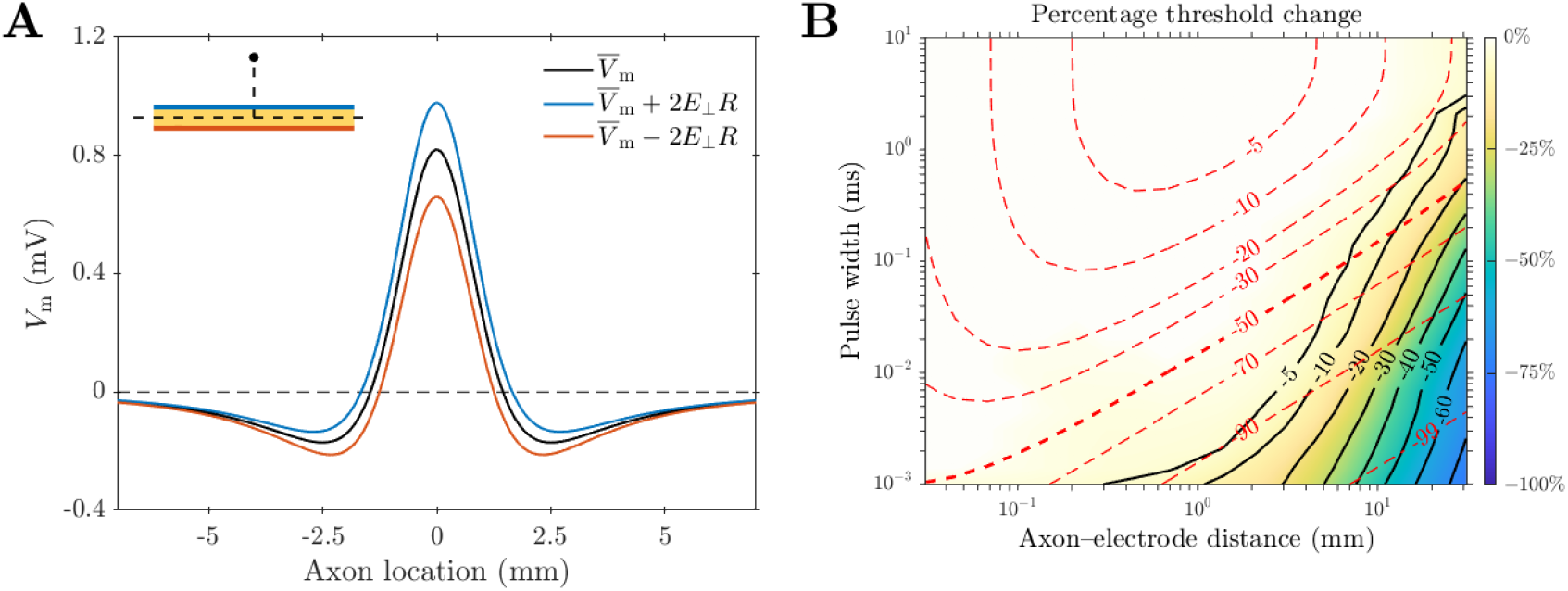
**A**. Illustration of transverse polarization of an axon near a point source cathode (inset) [8], showing the average transmembrane potential 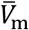 around the axon’s circumference as a function of axial position *x* (black), and transmembrane potentials along the cathodic (blue) and anodic (orange) sides perturbed by the transverse E-field *E*_⊥_ in opposite directions. Applying linear integrate-and-fire behavior to the pointwise transmembrane potential (peak of blue line) would result in lower thresholds for transverse polarization compared to the average polarization (peak of black line) used in the conventional cable equation. **B**. Relative difference for a combination of various axon-electrode distances and pulse durations (horizontal and vertical axes, respectively, both on logarithmic scales) comparing simulated thresholds of an unmyelinated 3 μm radius axon obtained with the modified cable equation that accounts for transverse polarization against thresholds obtained with the conventional cable equation, shown as black contour lines filled with colors in between. Transverse polarization has minimal effect on thresholds (light yellow region, less than −5% change), except for combinations of very large axon–electrode distances and short pulse durations not used in typical neural engineering applications (lower right corner). Overlaid red dashed contour lines show threshold changes predicated with linear integrate-and-fire behavior applied to the transmembrane potential pointwise, revealing extremely large discrepancies compared to the simulation results. Panel B is reproduced using data from [8].

Given the complexity of spatiotemporal interactions between the E-field and neurons, neural simulations should always be considered for determining activation thresholds. Simulations with the microscopic E-field [5] that were used to derive the statistical method [4] demonstrated that the activation thresholds were largely unaffected by microscopic E-field perturbations [5], directly contradicting the results by Weise et al. [4]. Therefore, microscopic E-field perturbations are unlikely to be the cause of TMS threshold discrepancy between computational models and *in vivo* measurements.

## Acknowledgments

This work was supported by the National Institute of Mental Health of the US National Institutes of Health under grant No. R01MH128422. The content is solely the responsibility of the authors and does not necessarily represent the official views of the funding agency. Computational support was provided by the Duke Compute Cluster.

## CRediT authorship contribution statement

B. Wang: Conceptualization, Investigation, Methodology, Formal analysis, Software, Visualization, Writing–original draft, Writing–review & editing. M. A. Hussain: Investigation, Writing–review & editing. T. Worbs: Investigation, Writing–review & editing. A. Thielscher: Conceptualization, Funding acquisition, Writing–review & editing. W. M. Grill: Conceptualization, Resources, Funding acquisition, Writing–review & editing. A. V. Peterchev: Conceptualization, Funding acquisition, Writing–review & editing.

## Footnotes

1 Here, microscopic refers to spatial scales on the order of tens of nm up to a few tens of μm. Any tissue structures on the order of 100 μm and above can no longer be homogenized on the macroscopic scale of the finite element models, e.g., 1 mm resolution of imaging data.

2 The secondary E-field ***E***_micro_ has zero mean not only in the direction aligned with ***E***_macro_ but also in other directions [5]. The zero mean of ***E***_micro_ is due to the conservation of charge generated on the membranes by the E-field, resulting in a multipolar distribution with a net zero charge [8].

3 Conservative potentials are used here for simplicity, whereas for the non-conservative E-field of TMS, the corresponding pseudo-potentials [10] should be used, following the same derivation.

4 Spatial variations on even larger scale are no longer microscopic and could be captured by mesoscopic changes in effective conductivity or by explicitly modeling their sources.

5 Specifically, the integration of the microscopic term in (1) is over the internodal distance, typically on the order of 1 mm, and therefore averages out to zero. Whereas for (2), the transmembrane potential 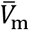 on the order of tens of mV dominates over *V*_e,micro_, which is at most on the order of 1 mV (assuming a very high microscopic E-field of 100 V/m and very large spatial wavelength of 50 μm, the maximum *V*_e,micro_ is 100 V/m × 50 μm /(2π) ≈ 0.8 mV).

